# Higher genetic risk of schizophrenia is associated with lower cognitive performance in healthy individuals

**DOI:** 10.1101/103622

**Authors:** Rebecca Shafee, Pranav Nanda, Jaya L. Padmanabhan, Neeraj Tandon, Ney Alliey-Rodriguez, Richard S. E. Keefe, Scot K. Hill, Jeffrey R. Bishop, Brett A. Clementz, Carol A. Tamminga, Elliot S. Gershon, Godfrey D. Pearlson, Matcheri S. Keshavan, John A. Sweeney, Elise B. Robinson, Steven A. McCarroll

**Author notes:** Corresponding Author: Rebecca Shafee.

## Abstract

Psychotic disorders including schizophrenia are commonly accompanied by cognitive deficits. Recent studies have reported negative genetic correlations between schizophrenia and indicators of cognitive ability such as general intelligence and processing speed. Here we compare the effect of the genetic risk of schizophrenia (PRS_SCZ_) on measures that differ in their relationships with psychosis onset: a measure of current cognitive abilities (the Brief Assessment of Cognition in Schizophrenia, BACS) that is greatly reduced in psychosis patients; a measure of premorbid intelligence that is minimally affected by psychosis (the Wide-Range Achievement Test, WRAT); and educational attainment (EY), which covaries with both BACS and WRAT. Using genome-wide SNP data from 314 psychotic and 423 healthy research participants in the Bipolar-Schizophrenia Network for Intermediate Phenotypes (B-SNIP) Consortium, we investigated the association of PRS_SCZ_ with BACS, WRAT and EY. Among apparently healthy individuals, greater genetic risk for schizophrenia (PRS_SCZ_) was associated with lower BACS scores (r = −0.19, p = 1 × 10^−4^ at P_T_ = 1 × 10^−4^) but did not associate with WRAT or EY, suggesting that these areas of cognition vary in their etiologic relationships with schizophrenia. Among individuals with psychosis, PRS_SCZ_ did not associate with variation in cognitive performance. These findings suggest that the same cognitive abilities that are disrupted in psychotic disorders are also associated with schizophrenia genetic risk in the general population. Specific cognitive phenotypes, independent of education or general intelligence, could be more deeply studied for insight into the specific processes affected by the genetic influences on psychosis.

**Significance:** Psychotic disorders such as schizophrenia often involve profound cognitive deficits, the genetic underpinnings of which remain to be elucidated. Poor educational performance early in life is a well-known risk factor for future psychotic illness, potentially reflecting either shared genetic influences or other risk factors that are epidemiologically correlated. Here we show that, in apparently healthy individuals, common genetic risk factors for schizophrenia associate with lower performance in areas of cognition that are impaired in psychotic disorders but do not associate independently with educational attainment or more general measures of intelligence. These results suggest that specific cognitive phenotypes – independent of education or general intelligence – could be more deeply studied for insight into the processes affected by the genetic influences on psychosis.

## Introduction

Schizophrenia is a debilitating psychiatric disorder that commonly involves severe cognitive deficits that compromise functional ability. Although at present schizophrenia is classified as a psychotic disorder, it has been suggested that it is primarily a cognitive disorder (1,2). Cognitive deficits have been reported not only in schizophrenia but also in other psychotic disorders (3,4). Underperformance in general intelligence tasks as well as tasks designed to be specific to cognitive domains such as memory, executive function, motor function etc. have been noted in psychosis patients (5).

Many of these cognitive deficits are present many years prior to the development of the illness (6,7). A meta-analysis of 4396 schizophrenia cases and 745,000 controls showed that every point decrease in premorbid IQ associates with a 3.7% increase in schizophrenia risk (8). In another analysis in a nationwide cohort of over 900,000 Swedish individuals, children with the lowest grades showed a 4-fold increased risk of developing schizophrenia and schizoaffective-disorder and a 3-fold increased risk of developing other psychotic illnesses (9). Additionally, studies of clinically high-risk (CHR) groups have shown that these near-psychotic individuals are cognitively impaired compared to healthy controls and that within the CHR group those that convert to psychosis in the future display lower cognitive performance compared to the nonconverters (10,11,12,13). There is also evidence of lower cognitive performance in first-episode psychosis patients compared to the CHR group (10,12,13). Together these results indicate that cognitive deficits are significantly associated with risk of developing a psychotic illness. It has also been reported that CHR individuals that convert to psychosis show longitudinal decline in neurocognitive performance as they progress to acute psychosis (14). Although the relative roles played by cognitive deficit and cognitive decline along the psychosis onset trajectory are still not well understood, it is clear that lower cognitive function is related to increased vulnerability to psychotic illness onset.

Both cognitive performance and schizophrenia are heritable (15–23), and significant genetic overlap has been reported between schizophrenia and indicators of cognitive ability, such as general intelligence or processing speed (24–30). However, it is still unclear how the genetic differences associated with schizophrenia influence cognitive function, and which domains of cognitive function are most associated with schizophrenia risk.

Motivated by these earlier findings, we investigated the relationship of the genetic risk for schizophrenia – as defined by large constellations of common variants that associate with schizophrenia risk (“polygenic risk”, PRS_SCZ_) – on three phenotypes: 1. the Brief Assessment of Cognition in Schizophrenia (BACS, 31), which consists of six subtests spanning multiple cognitive domains and provides a composite score of general cognitive function, 2. the Wide-Range Achievement Test (WRAT, 32–34) reading score, a measure of premorbid intellectual potential; and 3. educational attainment (as measured by years of education, EY), which is phenotypically associated with WRAT and BACS and also genetically overlaps with cognition (35,36). These analyses were done using the psychotic probands (PSYCH, N = 314) and nonpsychotic individuals (NPSYCH, N = 423) of the Bipolar-Schizophrenia Network of Intermediate Phenotypes consortium (B-SNIP, 3, 37). The PSYCH group included psychotic schizophrenia, bipolar disorder and schizoaffective disorder probands and the NPSYCH group was formed by combining healthy controls and nonpsychotic family members (more details in Methods).

These measures of cognitive performance and educational achievement vary in their relationships to schizophrenia. While psychosis probands show lower values for all three phenotypes compared to healthy control individuals, the deficit in BACS in the patient group is much more significant than the deficit in EY or WRAT (3). In our sample the psychotic patient group’s mean BACS performance was one standard deviation lower than the mean of the nonpsychotic group, whereas their EY and WRAT scores were lower by 0.3 standard deviations (Figures S1). Also, while many cognitive domains captured by BACS, such as, working memory and processing speed, show increasing deficits along the psychosis onset trajectory (decreasing performance from CHR individuals that do not convert to psychosis, to CHR individuals that do convert to psychosis to first-episode psychosis patients, 10-13), WRAT is a relatively stable measurement that is minimally affected by psychosis onset (10) and is commonly used as a measure for premorbid intelligence (32-34). Based on these phenotypic observations we hypothesized that the genetic risk of schizophrenia should be more strongly associated with BACS than with WRAT. Additionally, we investigated the effect of the polygenic predictors of educational attainment (PRS_EDUC_) on BACS, EY and WRAT. We find that in healthy individuals, genetic risk for schizophrenia is associated with BACS, but not with WRAT or EY. This finding suggests that specific domains of cognition may be more closely etiologically linked to schizophrenia than other domains are, creating an opportunity for longitudinal studies to identify the domains that best predict illness onset.

## Results

### Validation of genetic data: genetic risk for schizophrenia is higher among psychosis patients

Polygenic liability for a trait – an estimate of genetic risk – is calculated by adding the relatively small effects of many individual common variants, as estimated from very large earlier genetic studies (the 2014 schizophrenia Genome-Wide Association Study by PGC consisted of 36,989 cases and 113,075 controls, 22). The score can be used to predict the value of a continuous trait (e.g. height) or the risk of a disease. For schizophrenia, polygenic scores predict 18.4% of the variation in disease risk (22). These scores are often calculated at multiple p-value thresholds (P_T_). Lower p-value thresholds include fewer SNPs, each with more significant associations. Higher p-value thresholds include more SNPs, but the effect of the risk-conferring SNPs can be diluted among a larger fraction of neutral SNPs. This varying P_T_ approach is exploratory and used when the P_T_ for which the predictive power of the score is maximum for a particular phenotype is unknown.

An individual’s PRS_SCZ_ represents common genetic influences on the risk of developing schizophrenia. Hence one would expect higher PRS_SCZ_ in the PSYCH group compared to the NPSYCH group. As a validation analysis of our genetic data we investigated the differences in PRS_SCZ_ between the PSYCH and NPSYCH groups. As shown in Figure 1, compared to the NPSYCH group the PSYCH group showed significantly higher PRS_SCZ_ at all P_T_ as expected (p ≤ P_FDR_ = 8.9 × 10^−4^). Among the psychosis probands schizophrenia patients had highest PRS_SCZ_ (Figure S2). In our sample PRS_EDUC_ did not differ significantly between the PSYCH and the NPSYCH groups at any P_T_. Effect sizes and p-values for PSYCH/NPSYCH group differences in PRS_SCZ_ and PRS_EDUC_ can be found in Table S1. Figure S2 shows the distributions of PRS_SCZ_ and PRS_EDUC_ for the different DSM diagnosis groups.

**Figure 1:**
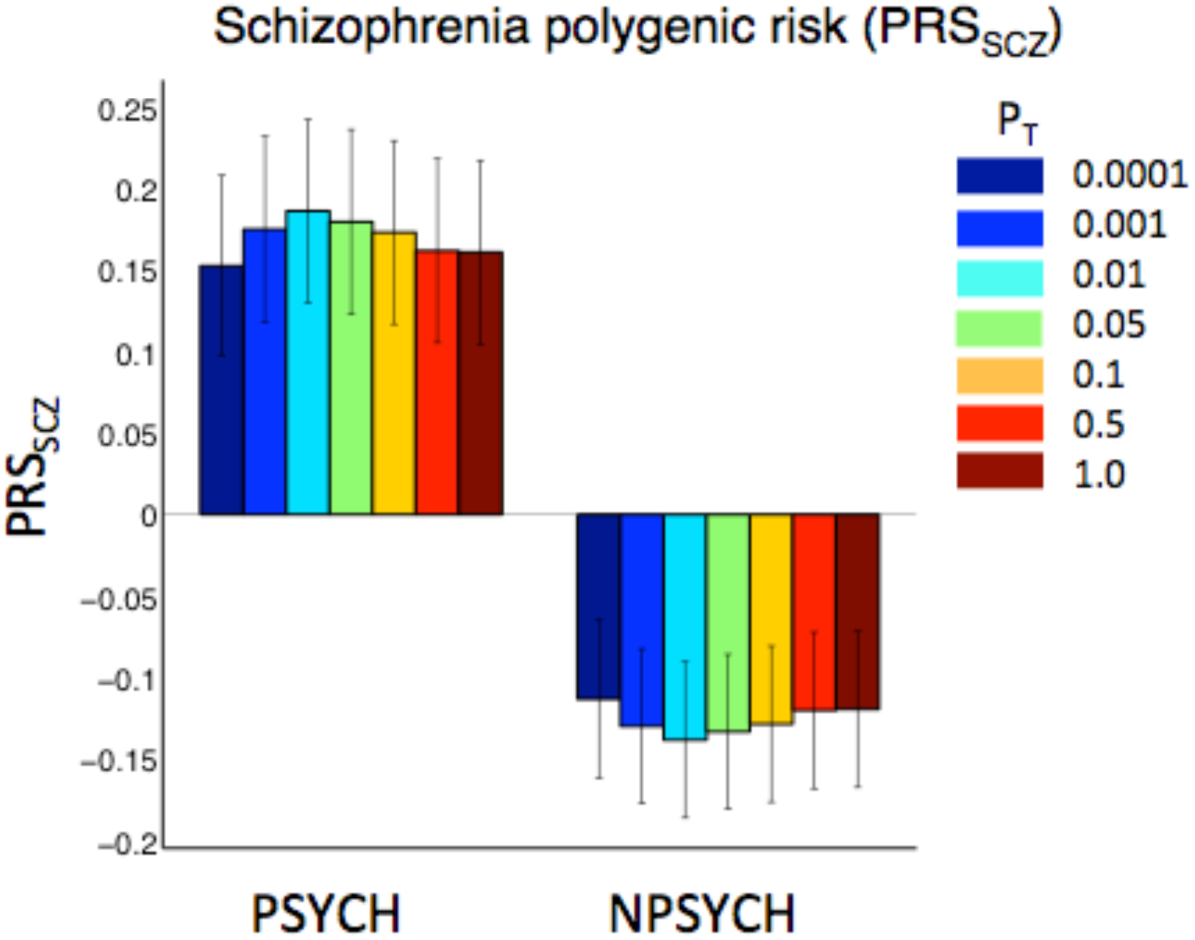
Mean Polygenic scores of schizophrenia (PRS_SCZ_) in the psychotic (PSYCH, N = 314) and the nonpsychotic (NPSYCH, N = 423) groups. The vertical black lines show the standard errors of the mean (SEM). Scores were calculated at seven p-value thresholds (P_T_): 0.0001, 0.001, 0.01, 0.05, 0.1, 0.5 and 1.0, which are shown in different colors. All scores were z-transformed before mean and SEM calculation. PRS_SCZ_ was significantly higher (p ≤ P_FDR_ = 8.9 × 10^−4^) in the PSYCH group compared to the NPSYCH group at all P_T_. Table S1 shows the p-values for this analysis.

### EY, BACS and WRAT are positively related as expected; psychosis does not alter these correlations

An individual’s educational attainment, cognitive functioning and intellectual potential are expected to be correlated with one another. The relationships among these three traits were similar in the two groups (Figure 2), indicating that the presence of psychosis did not alter the interdependence of these three phenotypes. Although BACS, WRAT and EY were significantly lower in the PSYCH group compared to the NPSYCH group, the effect size of deficit in BACS was three times as much as the deficits in EY or WRAT (BACS group difference Cohen’s d = −1.05, p = 2.3 × 10^−37^, Figure S1). Additionally, partial correlation analyses between pairs of these three phenotypes controlling for the third phenotype revealed that, 1. EY and WRAT shared a positive correlation that could not be accounted for by BACS; 2. WRAT and BACS shared a positive correlation that could not be accounted for by EY; and 3. although EY and BACS were weakly positively correlated, this correlation was mediated via factors that could be captured by WRAT (Table S2).

**Figure 2:**
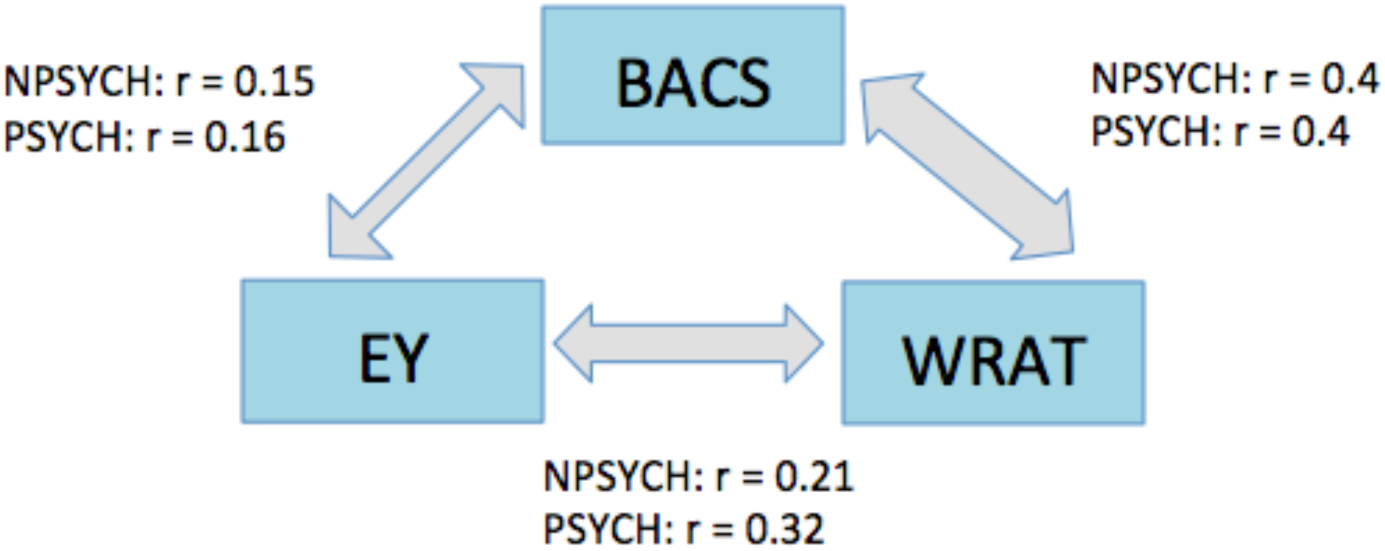
Relationship between BACS, educational attainment EY and WRAT. The three phenotypes were positively correlated in both the PSYCH (N = 314) and the NPSYCH (N = 423) groups, with the strongest correlation between BACS and WRAT (r ~ 0.4, p < 10^−11^). The magnitudes of the correlations were not significantly different between the psychotic (PSYCH) and the nonpsychotic (NPSYCH) groups. More details can be found in Table S2.

### Higher genetic risk of schizophrenia is associated with lower BACS score in nonpsychotic individuals

To evaluate whether the genetic risk of schizophrenia associates with variations in BACS, EY and WRAT, correlations of PRS_SCZ_ with these measures were calculated within the PSYCH and the NPSYCH groups separately. Since EY, BACS and WRAT were positively correlated (Figure 2), full correlations as well as partial correlations controlling for the remaining two traits were calculated. Figure 3 shows the strongest full and partial correlations for each phenotype. For all polygenic score correlation analyses the combined FDR corrected p-value threshold was determined to be P_FDR-PRS_ = 0.0064 (Methods). The numerical values for the correlation coefficients and the p-values for both groups at all P_T_ can be found in Table S3.

**Figure 3:**
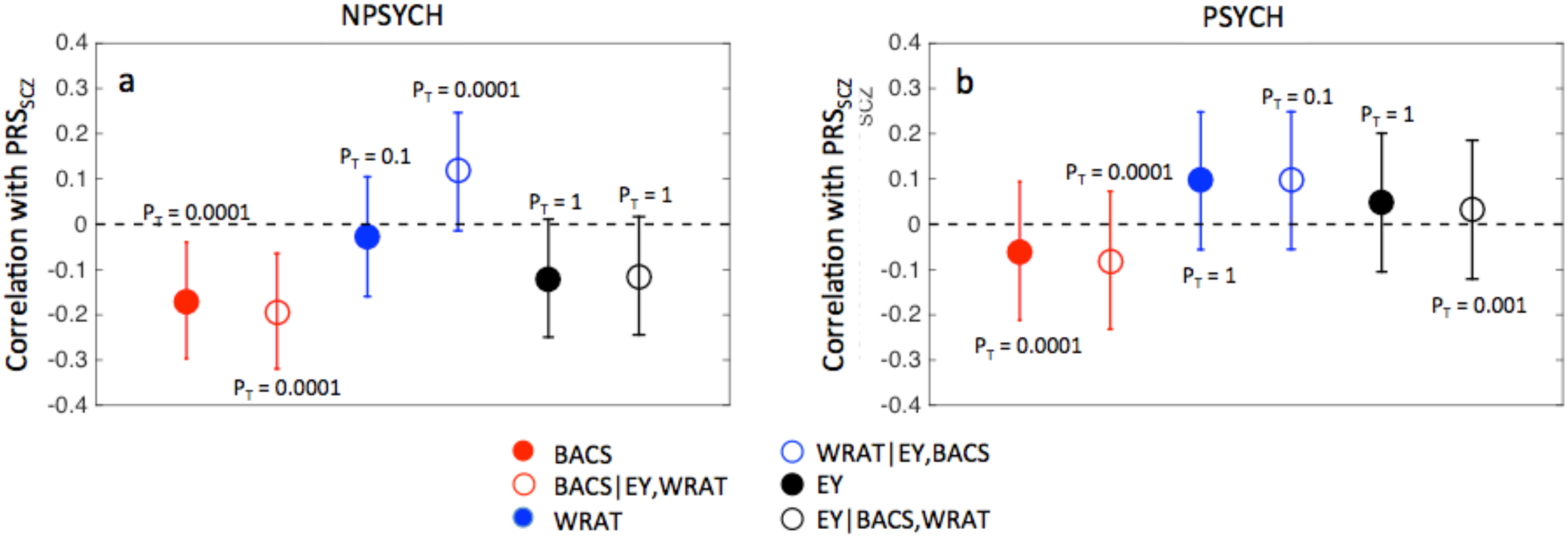
Correlations of polygenic score of schizophrenia (PRS_SCZ_) with BACS, WRAT and EY in the nonpsychotic (panel a, NPSYCH, N = 423) and the psychotic groups (panel b, PSYCH, N = 314). Only the strongest correlation for each phenotype is shown with the corresponding P_T_ labeled. Correlation coefficients are plotted with confidence intervals reflecting the FDR-corrected significance p-value of P_FDR-PRS_ = 0.0064. Confidence intervals not including the r = 0 line correspond to significant correlations. The effects of age, sex, collection site, ancestry principal components, DSM diagnoses (in the PSYCH group) and healthy control (HC)/ nonpsychotic family member (NPFAM) status (within the NPSYCH group) were regressed out of EY, BACS, WRAT and PRS_SCZ_. Within the NPFAM group, the respective proband’s DSM diagnosis was also added as a covariate. Full correlations and partial correlations (in each case controlling for the other two traits) with PRS_SCZ_ are shown. Correlation coefficients and corresponding p-values for all P_T_ can be found in Table S3.

As shown in Figure 3, PRS_SCZ_ showed insignificant correlations with WRAT and EY in both the PSYCH and the NPSYCH groups. Significant negative correlations were observed between PRS_SCZ_ and BACS in the NPSYCH group but not in the PSYCH group at multiple P_T_ (Table S3). The strongest correlation, as shown in Figure 3a, was observed when EY and WRAT were regressed out at P_T_ = 1 × 10^−4^ (r = −0.19 and p = 1.0 × 10^−4^). Since the NPSYCH group was formed by combining the healthy control (HC) and the nonpsychotic family member (NPFAM) subgroups (Methods), HC/NPFAM status was accordingly used as a covariate in the above correlation analyses. Additionally, the relationship between PRS_SCZ_ and BACS was explored within the HC and the NPFAM subgroups separately (supplementary material) to ensure these results did not arise as an artifact of merging the two subgroups. There was no significant difference in the correlation coefficients in the two subgroups (Table S4). At the subgroup level, statistically significant correlation between BACS and PRS_SCZ_ was seen at P_T_ = 10^−4^ in the HC group (r = −0.25, p = 1.9 × 10^−3^), which remained significant when EY and WRAT were regressed out. In the NPFAM subgroup, significant negative correlation was detected at P_T_ = 0.01 (r = − 0.19, p = 6.4 × 10^−3^) when EY and WRAT were regressed out (Table S4).

The PSYCH group consisted of schizophrenia, bipolar disorder and schizoaffective disorder probands. Correlation between BACS and PRS_SCZ_ within the schizophrenia probands only (SZP, N = 100) was also insignificant. Adding illness duration, number of hospitalization, chlorpromazine dose, number of psychotropic drugs, and social-functional scale score as covariates in the correlation analysis between BACS and PRS_SCZ_ did not alter the lack of significant results in the PSYCH group.

### Higher genetic score of educational attainment is associated with higher EY and WRAT

To investigate the effects of the common genetic polygenic influences on educational attainment (36) on these traits, we analyzed the correlation of the polygenic score of educational attainment (PRS_EDUC_) with EY, WRAT and BACS. As expected, PRS_EDUC_ showed significant positive correlations with EY in both the PSYCH group (Fig. 4b, strongest correlation of r = 0.19, p = 0.0016 at P_T_ =0.05) and the NPSYCH group (Fig. 4a, strongest correlation of r = 0.17, p = 7 × 10^−4^ at P_T_ = 0.01). When BACS and WRAT were controlled for, these significant correlations decreased and the only remaining significant correlation was in the NPSYCH group at P_T_ = 0.01 (r = 0.15, p = 0.003). Statistically significant positive correlations were observed between PRS_EDUC_ and WRAT in both the PSYCH group and the NPSYCH group (Figs. 4a,b) at several thresholds (strongest correlation of r = 0.26, p = 10^−5^ at P_T_ = 0.05 in PSYCH and strongest correlation of r = 0.15, p = 2.4 ×10^−3^ at P_T_ = 0.05 in NPSYCH). No significant correlation was found between PRS_EDUC_ and BACS in either group. The numerical values for all the correlation coefficients and p-values can be found in Table S5. Together these results indicate that the common variants associated with educational attainment influence education and premorbid intellectual potential (as measured by WRAT) more significantly than general cognitive performance in the general population.

## Discussion

Educational attainment (EY), premorbid intellectual potential (WRAT), and cognitive performance (BACS) are interrelated and correlated phenotypes (Figure 2), making it challenging and important to disentangle the effects of genetic risk on each. In the current work, in which we sought to disentangle these effects through simultaneous conditional analyses, the genetic risk of schizophrenia (PRS_SCZ_) significantly associated only with BACS and not with EY or WRAT in nonpsychotic individuals. PRS_SCZ_ explained about 4% of the variance in the composite BACS scores in the NPSYCH group.

These results may help to understand earlier results on genetic correlations of schizophrenia with other phenotypes. Using the UK Biobank sample Hagenaars et al. (29) found significant negative relationships between schizophrenia genetic risk and three cognitive traits: verbal-numerical reasoning, memory and reaction time. While the composite BACS score is not directly comparable to these three phenotypes, the six subtests used to calculate the BACS composite score significantly overlap with these three cognitive subdomains (Methods). Hence, our findings are consistent with the results in Hagenaars et al. We further found that the effect of the genetic risk of schizophrenia on BACS was significant after accounting for variability due to EY and WRAT. While BACS measures an individual’s ability to use cognitive resources to solve problems, WRAT is more of a measure of crystallized verbal knowledge. These results indicate that cognitive domains measured by BACS– rather than other brain phenotypes that shape premorbid intelligence or educational attainment – are likely more direct targets of the genetic risk factors of schizophrenia

Though we observed a strong negative correlation between PRS_SCZ_ and BACS at multiple P_T_ in the nonpsychotic group (NPSYCH, Figure 3), as well as individually within the healthy controls and the nonpsychotic family members (Table S4), we did not observe such a correlation among psychosis patients (PSYCH). Thus, we found no evidence that psychotic individuals at elevated genetic risk (due to common variants) are more severely cognitively impaired than psychotic individuals at lower genetic risk. Cognitive deficits in the patient group may thus reflect morbid factors that are not predicted by PRSscz. The morbid consequences of disease progression, protective effects of supportive care, and the effects of medications, medical and psychiatric comorbidity and substance use may have more profound effects on pathologic trajectories than any pre-existing variation in cognition, and may involve biological mechanisms distinct from those that mediated pre-onset risk. Recently, a similar result was reported in a study of Autism Spectrum Disorder (ASD) in which the polygenic risk of ASD did not predict IQ in the ASD probands (despite a strong positive correlation in the general population, 38) although the polygenic scores of educational attainment and schizophrenia did (39). This too could be due to non-genetic factors playing a significant role in determining cognitive function in the ASD patients, overshadowing the cognitive variability due to the genetic risk of developing the disease.

A higher value of PRS_EDUC_ reflects the inheritance of a larger-than-average number of genetic factors (alleles) that associate with educational attainment in the broader population. Therefore, the observed significant positive correlations between PRS_EDUC_ and EY in both the PSYCH and the NPSYCH groups in our sample were expected (Figure 4). Additionally, we found that PRS_EDUC_ was positively associated with WRAT in both the PSYCH and NPSYCH groups. The correlation between PRS_EDUC_ and BACS was not significant in either group. Together, these results imply that the common variants associated with educational attainment influence EY and WRAT more significantly than they influence BACS in the general population. It should be noted that Hagenaars et al. (29) also reported lack of significant correlations between their educational attainment polygenic score and memory and reaction time. However they did report significant positive correlation of education polygenic score with verbal-numerical reasoning, which is likely closer to our measure of WRAT than to BACS. In our sample of 737 individuals, the absence of correlation between two quantities does not necessarily imply the true absence of effects. Our study might be underpowered to detect weaker correlations (at P_FDR_=0.0064, 25% and 17% power to detect a correlation of 0.1 in the NPSYCH group and the PSYCH group, respectively).

**Figure 4:**
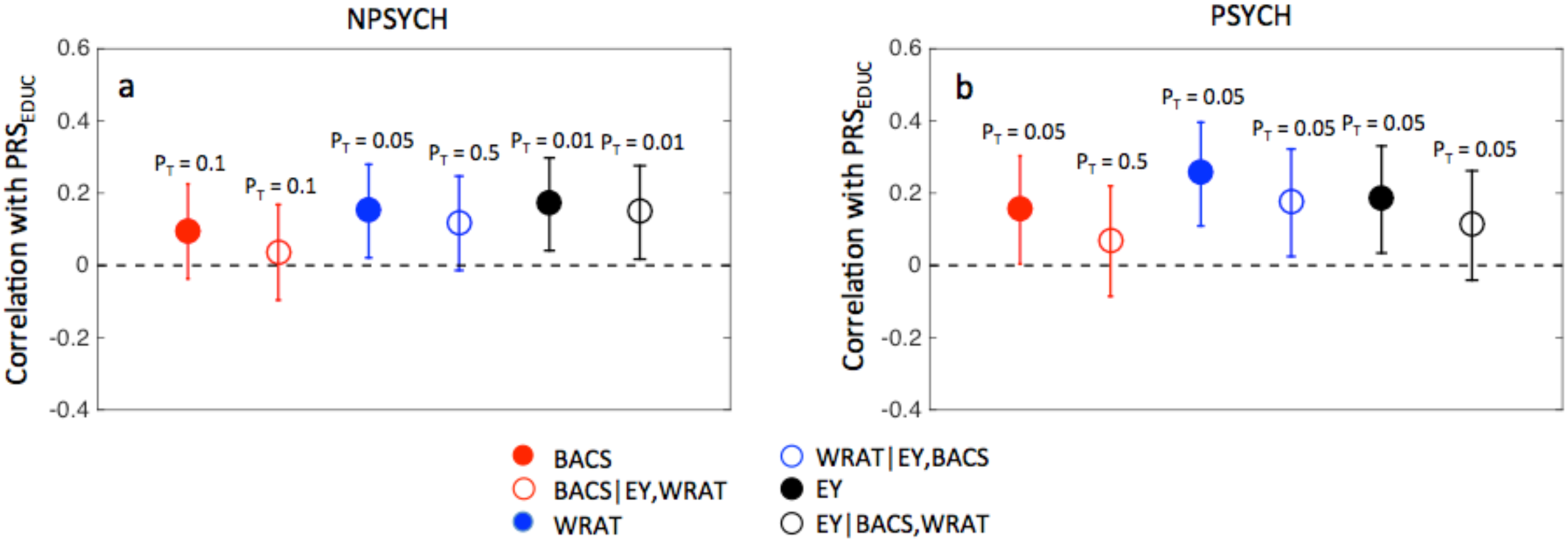
Correlations of polygenic score of educational attainment (PRS_EDUC_) with BACS, WRAT and EY in the nonpsychotic (panel a, NPSYCH, N = 423) and the psychotic groups (panel b, PSYCH, N = 314). Only the strongest correlation for each phenotype is shown with the corresponding P_T_ labeled. Correlation coefficients are plotted with confidence intervals reflecting the FDR-corrected significance p-value of P_FDR-PRS_ = 0.0064. Confidence intervals not including the r = 0 line correspond to significant correlations. The effects of age, sex, collection site, ancestry principal components, DSM diagnoses (in the PSYCH group), and healthy control (HC)/ nonpsychotic family member (NPFAM) status (in the NPSYCH group) were regressed out of EY, BACS, WRAT and PRS_SCZ_. Within the NPFAM group, the respective proband’s DSM diagnosis was also added as a covariate. Full correlations and partial correlations (in each case controlling for the other two traits) with PRS_EDUC_ are shown. Correlation coefficients and corresponding p-values for all P_T_ can be found in Table S5.

Decreased performances on all three cognitive measures used in this work: BACS, WRAT and EY, could be considered as risk factors for future psychotic illness. However, the phenotypic deficit in BACS in psychotic probands is much more significant compared to the deficits in EY and WRAT. Additionally, while the cognitive domains measured by BACS show increasing deficits along the psychosis onset trajectory during the high-risk prodromal phase, WRAT remains a stable measure during this period influenced primarily by events occurring in earlier life well before the illness prodrome. In this study, genetic risk of developing schizophrenia was more strongly connected to cognitive performance among unaffected individuals than to either (i) the magnitude of cognitive impairment among psychosis patients, or (ii) educational attainment or premorbid intellectual potential (as measured by WRAT reading scores). Our results indicate that BACS is genetically more closely associated with schizophrenia and possibly psychotic illnesses in general, compared to the other two measures in spite of the interrelated nature of these phenotypes. These results suggest that specific cognitive phenotypes, independent of education or general intelligence, could be more deeply studied for insight into the specific processes affected by the genetic influences on psychosis risk. Studies of the premorbid period and/or the high-risk prodromal stage may be particularly valuable for understanding the connections and sequences of events involving genetic risk, performance in specific cognitive domains, and chronic psychotic illness.

## Methods

### Study design and participants

This study includes 737 Caucasians (Table S6) from the Bipolar-Schizophrenia Network for Intermediate Phenotypes (B-SNIP), which is a multi-site consortium studying risk factors for psychosis (3, 37). Previous work using this cohort reported BACS performance to be consistent with a dimensional model of psychosis in which the schizophrenia probands were most severely impaired, followed by schizoaffective disorder probands and bipolar disorder probands (3). Also, a recent study using the B-SNIP cohort reported the lack of a biological basis for the segmenting of the psychoses by DSM diagnoses (40). Due to these findings and also due to the relatively small sample size we combined all psychotic probands to form the PSYCH group (N = 314). To control for effects that might arise from differences among the schizophrenia/bipolar/schizoaffective disorder probands, the DSM diagnosis of all probands was used as a covariate for correlation analyses within the PSYCH group. The NPSYCH group consisted of nonpsychotic individuals and was created by combining healthy controls (HC, N = 180) and first-degree relatives with no history of psychosis (NPFAM, N = 243). It was shown in (3) that nonpsychotic relatives with elevated axis II traits (cluster A or cluster B) were significantly cognitively impaired compared to healthy controls. To minimize heterogeneity within the NPSYCH group, these individuals were excluded. Additionally, HC/NPFAM status as well as the respective proband’s DSM diagnosis (for NPFAM samples only) were used as covariates for all correlation analyses within the NPSYCH group. In the families for which multiple family members’ data were available, only one member selected at random from each family was included in the NPSYCH group.

Details of participant selection criteria can be found in previous publications from the B-SNIP consortium (3). Institutional review boards at each site approved the study and all sites used identical diagnostic, clinical, and recruitment techniques (37).

### Cognitive measures

Cognitive function was measured by the Brief Assessment of Cognition in Schizophrenia (BACS), which is a 30-minute test of global neuropsychological function (31). The BACS consisted of six subtests: verbal memory (VM), digit sequencing (DS), verbal fluency (VF), token motor task (TM), digit symbol coding (DSC) and Tower of London (TL). These tests encompass four cognitive domains: verbal memory, processing speed, reasoning and problem solving, and working memory (3). Premorbid intellectual potential was measured using the reading score of the Wide Range Achievement Test (WRAT IV), which has a phenotypic correlation of ~ 0.4 with full-scale intelligent quotient (32, 41). Self-reported years of education completed at the time of recruitment was used as a measure of educational attainment (EY).

### Collection and quality control of genetic data

Genetic data for the B-SNIP project were collected for 2053 subjects (multi-ethnic sample) using the Illumina Infinium PsychArray BeadChip™ platform. Genotypes underwent quality control using PLINK 1.9 (42, 43) based on a standardized protocol (44) in which individual markers were removed if they had a missing rate greater than 5%, deviated from Hardy-Weinberg equilibrium (p < 10^−6^), had a very low minor allele frequency (< 0.01), or demonstrated significantly different call rates between psychiatric probands and controls (p < 10^−5^). Subjects were removed for discordant sex information, outlying heterozygosity (> 3 standard deviations above the mean), or excessive missing genotype data (> 0.1). Kinship analyses were run using both PREST-plus (45) and KING (46) to confirm within-family kinships as well as to check for cryptic relatedness between unrelated individuals. Samples showing a relationship closer than 3rd degree to another unrelated individual were excluded. Samples of relatives that failed or showed a misrepresented kinship, or monozygotic twins were also excluded. After these initial quality control steps 1962 multi-ethnic samples remained.

### Ancestry Verification and Final Sample

Only samples with nonmissing age, sex, data collection site, BACS, WRAT, education, and history of psychosis were retained resulting in 1528 samples of whom 927 were self reported Caucasians (SRC). To avoid population stratification only SRC samples were used in all analyses. The ancestries of these SRC samples were verified by principal component analysis combining the B-SNIP genotype data with the 1000 Genomes phase 1 data (47). Samples that were more than four standard deviations away from the SRC group mean along the first ten principal components were excluded (N = 49). In the remaining sample only one first-degree relative of each proband was kept at random in order to avoid having related individuals in the NPSYCH group. Additionally, relatives with elevated cluster A or cluster B traits were removed. This final B-SNIP sample (N = 737) along with the different ethnic groups of the 1000 genome sample can be seen in the principal component scatter plot in Figure S4. The first ten principal components (PCs) of the ancestry analysis were used as covariates for all genetic correlation analyses to reduce the effects of genetic variability due to ancestry.

### Imputation of genotyped data and polygenic score construction

Imputation of genetic data was performed using HAPI-UR for pre-phasing (48) and IMPUTE2 for imputation (49,50) using a multiethnic reference panel (51). Chromosomes were phased separately, and then divided into 5-million-base-pair chunks for imputation. The 1000 Genomes phase 1 data were used as a reference panel for imputation (47). Poorly imputed single nucleotide polymorphisms (SNPs) were filtered post-imputation by removing SNPs that had information score less than 0.5 (52).

For a given set of SNPs, polygenic risk score was calculated for an individual as a weighted sum of that individual’s number of risk alleles at each SNP in the SNP set, with each allele weighted by the log of the odds ratios from the discovery sample, using custom scripts. Schizophrenia polygenic profile scores (PRS_SCZ_) and educational polygenic score (PRS_EDUC_) were calculated using the summary statistics of the Psychiatric Genome Consortium (PGC) schizophrenia GWAS meta-analysis results (22) and the summary statistics from Okbay et al. (36), respectively. Both scores were calculated for seven p-value thresholds of significance of association: P_T_ < 10^−4^, 0.001, 0.01, 0.05, 0.1, 0.5 and 1.0. Multiple values of P_T_ were investigated instead of a single threshold as the correlation analyses between the genetic profile scores and the cognition traits were exploratory. Scores calculated using higher P_T_ include more SNPs that are less significantly associated with the phenotype.

### Statistical Analysis

All statistical analyses were performed using Matlab (2012b). Group differences were calculated after regressing out the effects of age, sex, data collection site and the first ten principal components from the genetic ancestry analysis. The group difference p-values were calculated using the non-parametric Kruskal-Wallis test and effect sizes were calculated using the standard pooled variance Cohen’s d approach. Partial correlations between two variables while controlling for other variables were calculated using the Spearman Rank method (“partialcorr” function in Matlab) to minimize unwanted effects of outliers. Age, sex, data collection site, the first 10 principal components from the genetic ancestry analysis, and DSM diagnosis (schizophrenia/bipolar disorder/schizoaffective disorder status for members of the PSYCH group and respective relative’s diagnosis for member’s of the NPFAM group) were regressed out for correlation analyses within each group. As an additional precaution, the samples’ HC/NPFAM status was used as a covariate for all analyses within the NPSYCH group.

To correct for multiple hypotheses testing a false discovery rate (FDR) approach was used (53) instead of the more severe Bonferroni correction (54) as many of the variables, including polygenic scores calculated at different P_T,_ were correlated. Analysis-specific FDR thresholds as well as a threshold combining all p-values in this work were calculated at level α = 0.05. For the entire study P_FDR-ALL_ was 0.011. For analyses with polygenic scores the combined P_FDR-PRS_ was 0.0064 (Table S3, Table S4, Table S5). For group difference analyses in BACS, EY and WRAT between the PSYCH and the NPSYCH groups (Fig S1) and between the HC and the NPFAM groups (Fig S3) P_FDR_ was 0.022. P_FDR_ for group differences in PRS_SCZ_ and PRS_EDUC_ was 0.00089 (Table S1). For correlation analysis between EY, BACS, and WRAT (Table S2) P_FDR_ was 0.006. Significant results are reported based on analysis-specific FDR p-values since in most cases these values were smaller than P_FDR-ALL_. Relevant significance thresholds are mentioned with each result.

Our sample size was not sufficiently large to implement the recently developed statistical genetics methods of Linkage Disequilibrium (LD) Score regression (55) and Genome-Wide Complex Trait Analysis (GCTA, 56), so we focused on the polygenic score approach.

